# A Structure-Based Computational Pipeline for Broad-Spectrum Antiviral Discovery

**DOI:** 10.1101/2025.07.29.667267

**Authors:** Maria A Castellanos, Alexander M Payne, Jenke Scheen, Hugo MacDermott-Opeskin, Iván Pulido, Blake H Balcomb, Ed J Griffen, Daren Fearon, Haim Barr, Noa Lahav, David Cousins, Jessica Stacey, Ralph Robinson, Bruce Lefker, John D Chodera

## Abstract

The rapid emergence of viruses with pandemic potential continues to pose a threat to public health worldwide. With the typical drug discovery pipeline taking an average of 5–10 years to reach clinical readiness, there is an urgent need for strategies to develop broad-spectrum antivirals that can target multiple viral family members and variants of concern. We present a structure-based computational pipeline designed to identify and evaluate broad-spectrum inhibitors across viral family members for a given target in order to support spectrum breadth assessment and prioritization in lead optimization programs. This pipeline comprises three key steps: (1) an automated search to identify viral sequences related to a specified target construct, (2) pose prediction leveraging any available structural data, and (3) scoring of protein-ligand complexes to estimate antiviral activity breadth. The pipeline is implemented using the `drugforge` package: an open-source toolkit for structure-based antiviral discovery. To validate this framework, we retrospectively evaluated two overlapping datasets of ligands bound to the SARS-CoV-2 and MERS-CoV main protease (M^pro^), observing useful predictive power with respect to experimental binding affinities. Additionally, we screened known SARS-CoV-2 M^pro^ inhibitors against a panel of human and non-human coronaviruses, demonstrating the potential of this approach to assess broad-spectrum antiviral activity. Our computational strategy aims to accelerate the identification of antiviral therapies for current and emerging viruses with pandemic potential, contributing to global preparedness for future outbreaks.

## Introduction

Viral infections remain a persistent global threat, as demonstrated by the COVID-19 pandemic, during which the rapid spread of SARS-CoV-2 resulted in approximately 2 million deaths before vaccines became widely available (***Boby et al., 2023***; ***Carvalho et al., 2021***). Several factors, including human-driven biodiversity loss and climate change, increase the risk of emerging and re-emerging viral threats (***Vidal, 2020***; ***von Delft et al., 2023***), while globalization and high human mobility accelerate their potential to spark pandemics in the near future (***Goldin, 2014***). Despite these growing risks, antiviral discovery remains largely underfunded (***von Delft et al., 2023***), and mainly focused on developing inhibitors for individual viral protein targets to minimize interference with host cellular functions (***Geraghty et al., 2021***; ***Adalja and Inglesby, 2019***). This target-specific approach limits the development of new therapeutics in a landscape where novel viral threats can emerge in a matter of months. A more effective strategy is to design antivirals that target multiple viral family members by leveraging conserved structural features, enabling a faster and more adaptable response to future outbreaks (***Naqvi et al., 2020***; ***Schake et al., 2023***).

Antiviral drug development traditionally follows one of two main strategies: (1) targeting host proteins involved in the viral life cycle or (2) directly inhibiting essential viral proteins. While hosttargeted therapies offer the potential for broad-spectrum activity, they also carry a higher risk of toxicity due to interference with essential cellular processes (***Geraghty et al., 2021***). Furthermore, the intense forward selection on host proteins that are required for viral growth (***Wang et al., 2020***) or affect innate immune responses (***Enard et al., 2016***; ***Judd et al., 2021***), makes the efficacy of antivirals that modify such targets vulnerable to human variation. A more selective approach to broad-spectrum antiviral design would involve identifying conserved features among viral proteins, as evolutionary conservation often reflects functional significance within a viral family. These arguments are becoming increasingly persuasive as the amounts of sequence information increase (***Bloom and Neher, 2023***). In particular, proteins with conserved binding pockets are considered robust targets, as essential viral functions impose constraints on sequence variability, limiting the accumulation of mutations (***von Delft et al., 2023***; ***Liu et al., 2020***). Furthermore, even when mutations occur, key protein-ligand (P-L) interactions are often preserved, maintaining druggable features across viral variants. This study focuses on identifying and leveraging these conserved structural features to develop broad-spectrum inhibitors of viral proteins. Specifically, we target the main protease (M^pro^) of coronaviruses, an essential enzyme for viral replication (***Dai et al., 2020***). The conservation of M^pro^ across the coronavirus family makes it a compelling antiviral target for broad-spectrum inhibition (***von Delft et al., 2023***; ***Naqvi et al., 2020***).

Identifying conserved viral targets based on sequence similarity is a key step in broad-spectrum antiviral discovery. However, experimentally determined crystal structures for these related targets may not necessarily be available, making it difficult to apply the powerful tools of structure-based drug discovery in designing for the desired spectrum breadth. Recent advances in computational protein structure prediction now allow us to overcome this limitation, providing accurate structural models without the need for high-resolution crystallographic data. AI-driven protein folding models, such as AlphaFold2 (***Jumper et al., 2021***; ***Evans et al., 2021***) and RoseTTAFold (***Baek et al., 2021***), as well as newer approaches—including AlphaFold3 ***Abramson et al. (2024***), ESM-Fold (***Lin et al., 2023***), OpenFold ***Ahdritz et al. (2022***), and co-folding methods like Chai-1 (***Chai Discovery, 2024***) and Boltz (***Wohlwend et al., 2024***; ***Passaro et al., 2025***)—can predict the 3D co-ordinates of all heavy atoms in a protein by leveraging evolutionary information from multiple sequence alignments (MSA) and spatial constraints learned from structures in the Protein Data Bank (PDB) (***Jumper et al., 2021***; ***Mirdita et al., 2022***; ***Berman et al., 2003***). When modeling an evolutionarily conserved protein such as coronavirus M^pro^, incorporating an experimentally solved template structure can improve prediction accuracy while reducing computational cost (***Mirdita et al., 2022***; ***Baek et al., 2021***). In this study, we apply AlphaFold2 to a set of sequence-aligned coronavirus homologs, generating structural models based on a reference SARS-CoV-2 M^pro^ structure. These models serve as the foundation for structure-based drug design (SBDD) workflows, allowing us to systematically evaluate ligand binding across conserved targets and identify inhibitors with broad-spectrum potential.

Structure-based drug design is an essential component of modern drug discovery programs, leveraging 3D structural information of target proteins to predict and optimize protein-ligand (P-L) interactions (***Anderson, 2003***) to improve ligand potency. Given a bound pose, computational scoring functions estimate binding affinity based on electrostatic and steric interactions within the ligand pocket. These functions are typically classified as empirical (***Guedes et al., 2018***; ***Mcgann et al., 2003***; ***Trott and Olson, 2010***), knowledge-based (***Muegge and Martin, 1999***; ***Gohlke et al., 2000***), physics-based (also referred to as force field-based) (***Mey et al., 2020***), and, more recently, machine learning (ML)-based (***McNutt et al., 2021***; ***Liu and Wang, 2015***). While docking and scoring approaches are widely used to rank ligand poses, their low accuracy in predicting binding affinity across evolutionary variants remains a challenge (***Liu and Wang, 2015***). By integrating structural conservation principles with robust scoring functions, we aim to improve the prioritization of broad-spectrum inhibitors while keeping the computational cost low.

In this work, we present a study of broad-spectrum antiviral prediction with *spectrum*: a structure-based pipeline that integrates sequence alignment, protein structure prediction, docking, and scoring to systematically evaluate ligand binding across viral homologs. By leveraging scoring methods aimed at quantifying structural conservation, our approach aims to identify inhibitors with broad-spectrum potential. The pipeline is implemented as part of the `drugforge` package, an open-source toolkit based on the work by the ASAP Discovery Consortium to support modular, customizable workflows for antiviral drug discovery (see Section Data and Code Availability). The manuscript is divided as follows: we first introduce the sequence-to-structure pipeline, followed by a retrospective validation with experimental X-ray crystal structures and binding affinities of SARS-CoV-2 and MERS-CoV M^pro^. Finally, we apply our pipeline to a panel of coronaviruses with the antiviral Ensitrelvir, evaluating its accuracy in identifying broad-spectrum inhibitors.

## Results

### The AI-driven structure-enabled antiviral platform (ASAP) provides a test-bed for broad spectrum antiviral discovery

SARS-CoV-2 continues to pose a serious global health threat. The COVID-19 pandemic exposed major gaps in our ability to respond rapidly to emerging pathogens, including the lack of clinicaltrial-ready antivirals to address sudden outbreaks (***Boby et al., 2023***). Furthermore, the risk of future pandemics is ever-increasing (***Madhav et al., 2017***). To improve preparedness for future pandemics, it is critical to develop a globally equitable portfolio of therapeutics targeting viruses with high pandemic potential (***Griffen et al., 2024***). The AI-driven Structure-enabled Antiviral Platform (ASAP) AViDD Center is a multi-institutional consortium comprising over 140 academic and industry researchers, focused on accelerating antiviral discovery for high-risk viral families such as coronaviruses, flaviviruses, and picornaviruses (***ASAP Discovery Consortium, 2023***). As part of this consortium’s efforts, more than 1,100 crystal structures of SARS-CoV-2 and MERS-CoV main protease (M^pro^) have been elucidated via X-ray crystallography, and more than 3,300 ligand potency measurements have been gathered across these targets. In addition to generating large-scale experimental data, ASAP has developed a structure-enabled platform that integrates AI/ML and computational chemistry tools to support antiviral discovery campaigns for novel viral targets. In this study, we build on these efforts by leveraging the predictive capabilities of the ASAP computational infrastructure and the consortium’s experimental outputs to develop and validate a computational pipeline for identifying inhibitors with cross-variant activity within a viral family. We focus on the coronavirus family as a case study to evaluate the potential of this approach.

As part of the ASAP discovery consortium, we developed the `drugforge` package: an opensource, easy-to-install, MIT-licensed, fully integrated toolkit designed to support structure-based and ligand-based antiviral discovery (see Section Data and Code Availability). Built to streamline the structure-based drug discovery process, this package modularizes the computational drug discov-ery process, offering dedicated subpackages for key steps such as protein preparation, structure modeling, ligand pose prediction, molecular simulation, and binding-affinity scoring of P-L complexes. Each of these subpackages can be used independently or combined into customizable, end-to-end workflows that streamline lead discovery and optimization. In this study, we leverage the `drugforge` toolkit to build a pipeline focused on broad-spectrum antiviral discovery.

### Template-based AlphaFold followed by ligand transfer allows for pose and potency prediction of viral homologs

Broad-spectrum activity across a viral family can be associated with the conservation of key P-L interactions within the binding pocket (***von Delft et al., 2023***). Our approach to broad-spectrum antiviral discovery combines sequence alignment to identify viral homologs (i.e., proteins that share significant sequence similarity) with expert-informed filtering to prioritize targets with the highest potential for driving future human epidemics. The `drugforge-spectrum` package (see Data and Code Availability section) within the broader `drugforge` toolkit, implements this approach by providing a set of tools for 1) identifying viral homologs related to a given target protein (e.g., SARS-CoV-2 M^pro^), 2) generating P-L complex models with the assistance of available structural data, and 3) scoring ligand potency across homologs using a range of computational scoring methods.

The pipeline supports both the identification of viral homologs that may be susceptible to broad-spectrum inhibition and the evaluation of candidate compounds through a sequence-to-structure approach. Our strategy begins with a sequence-based search and alignment to identify relevant protein targets, followed by template-guided structure prediction using AlphaFold2. Ligand binding poses are then modeled, and P-L interactions are evaluated. By combining sequence-based search with AI-powered folding, this approach enables direct assessment of the relationship between sequence and structural conservation in the binding pocket, an essential factor for designing antivirals that retain activity across multiple related targets.

An overview of the pipeline is shown in Figure 1. The pipeline begins with a reference P-L crystal structure as input. In **step 1**, a BLAST search is performed to identify homologous proteins based on sequence similarity. The resulting list of sequences can then be filtered using some relevant criteria (e.g., viral host species). An example output of this step, filtering by targets that infect humans, is shown in Figure 2a. Next, the selected sequences are folded using the AlphaFold2-multimer model (***Evans et al., 2021***), as implemented in the open-source software ColabFold (***Mirdita et al., 2022***), with the original crystal structure used as a template (**step 2**). We choose the AlphaFold2-multimer model because it supports per-chain template inclusion, improving prediction accuracy at conserved binding sites. While other structure prediction tools could also be used (e.g., Open-Fold (***Ahdritz et al., 2022***), Boltz ***Passaro et al. (2025***)), AlphaFold2-multimer is available through ColabFold, which offers a fast, easy-to-install interface that integrates well with our existing infrastructure. The resulting protein models are then structurally aligned to the reference complex to ensure consistent binding pocket orientation. Using a template-based folding strategy preserves structural conservation with respect to the reference, and significantly reduces computational time (by approximately 10× compared to homology-search modeling, in addition to the 40–60× speed up offered by ColabFold compared to the standard AlphaFold2 implementation). Specifically, be-cause the multiple sequence alignment (MSA) is already generated during the BLAST step, it does not need to be recomputed during the AlphaFold modeling. Folded protein models for coronaviruses infecting humans, produced after this step, are presented in Figure 2b, aligned with SARS-CoV-2 in complex with Ensitrelvir (***Noske et al., 2023***).

**Figure 1.**
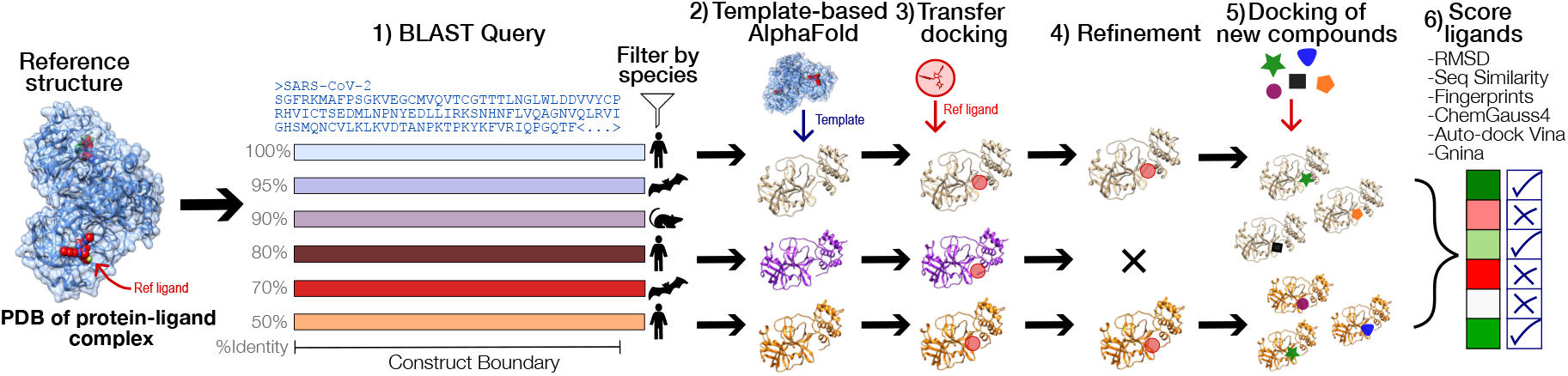
The affinity across a viral family can be predicted with a combination of sequence similarity search, docking, and affinity calculations. Starting from the PDB crystal structure of the reference protein and ligand complex (e.g., a SARS-CoV-2 M^pro^ dimer), a BLAST search is performed to find other proteins belonging to the same family. A few sequences of interest are selected given a filter criteria (e.g., all the viral proteins that infect humans) and then folded using the AlphaFold2-multimer model. Then, the coordinates of the reference ligand are transferred to the apo-protein, under the assumption that the binding pockets are similar. The new complex is docked, and P-L interactions are optimized with molecular mechanics force field minimization. The refined complex structures can then be used to dock new compounds and the inhibitors are scored by a combination of different methods.

**Figure 2.**
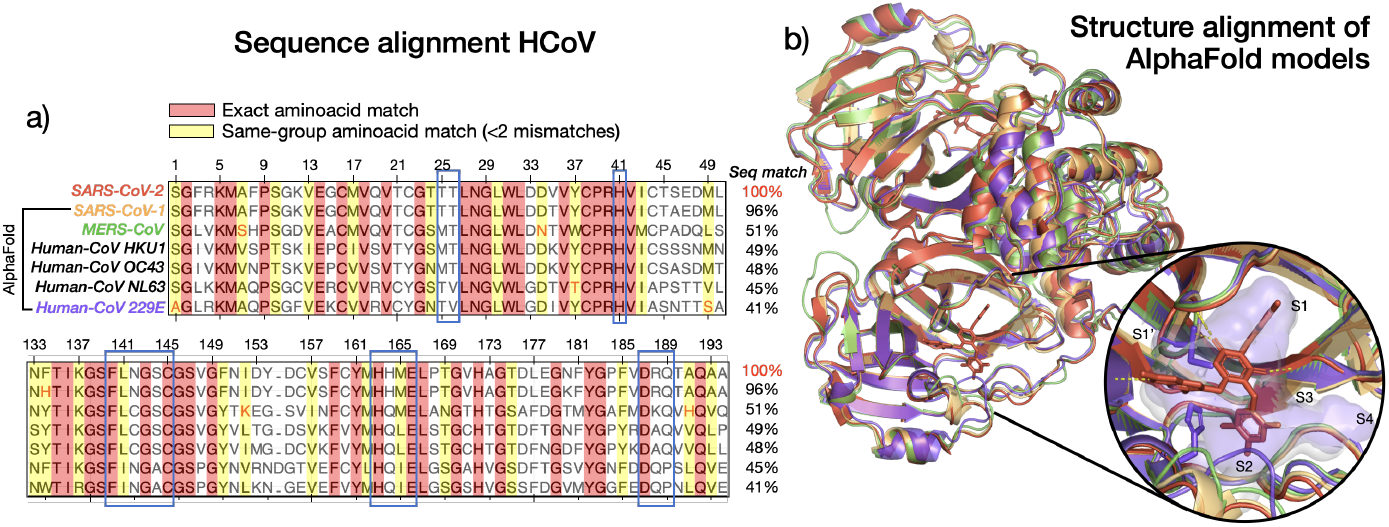
The M^pro^ binding pocket is highly conserved across the coronavirus family. (a) Multiple sequence alignment on proteins of the coronavirus family that infect humans, with exact aminoacid matches highlighted in red and same-group matches highlighted in yellow. The alignment shows significant conservation of the sequence, especially in the residues belonging to SARS-CoV-2 M^pro^ binding pocket (< 5A from ligand, outlined in blue). (b) Coronavirus proteins on the left are folded with the AlphaFold2 multimer model and aligned to SARS-CoV-2 M^pro^, shown here in complex with Ensitrelvir (8dz0 ***Noske et al. (2023***)), with each protein colored as indicated in a). The insert contains a zoomed view of the binding pocket with polar contacts as yellow dashed lines, as well as the pocket surface of the reference (white) and Human-CoV 229E (purple), with all pockets subunits indicated.

In **step 3**, the ligand coordinates from the reference structures are transferred to the prepared and aligned protein models, and docking is performed using the POSIT method (***Kelley et al., 2015***). The extrapolation of the reference ligand coordinates is only possible because the pipeline assumes the binding pocket of the viral family is evolutionary conserved. Minor sequence differences among homologs can introduce steric clashes in the transferred poses, which are resolved in **step 4** via molecular dynamics (MD) energy minimization. These refined P-L models can then be used for docking additional compounds (**step 5**), also using POSIT. Finally, in **step 6**, we predict the ligand affinities of the models prepared on either step 4 or 5 by using a variety of scoring techniques, enabling comparative evaluation of candidate compounds across viral homologs.

To evaluate ligand binding across multiple viral targets, we selected scoring functions that satisfy one or more of the following criteria: **1)** accurate prediction of binding affinity, **2)** reliable discrimination between active and inactive compounds and/or accurate ranking of candidate molecules, **3)** containing information on shared features across different protein-complexes, **4)** computational efficiency suitable for screening medium-to-large compound libraries, and **5)** availability as open-source software with straightforward installation. In this manuscript, we tested and evaluated scoring techniques spanning three major categories: **reference similarity-based, empirical**, and **machine learning (ML)-based** methods. For each category, we selected at least one representative technique to assess their individual and combined performance for broad-spectrum compound screening, based on the criteria above. The scoring methods and functions used are described in Table 1

**Table 1.** Scoring methods used to estimate protein-ligand binding of viral homologs

| Scoring Method | Scoring Function | Descriptors | Ref. |
| --- | --- | --- | --- |
| Reference similarity-based metrics | RMSD | Root-mean-square deviation of ligand atoms from reference pose | *Kabsch (1976)* |
|  | Sequence Similarity | Amino acid sequence identity between model and ref, for all residues and at the binding pocket | – |
|  | PLIF | Tversky similarity of P-L interaction fingerprints (by type and residue) | *Salentin et al. (2015); Adasme et al. (2021)* |
| Empirical scoring functions | ChemGauss4 | Gaussian-based shape complementarity, and hydrogen bonds with protein and with implicit solvent | *OpenEye, Cadence Molecular Sciences, Inc. (2025)* |
|  | AutoDock Vina | Steric fit, repulsion, hydrophobicity, and hydrogen bonding | *Trott and Olson (2010); Eberhardt et al. (2021)* |
| ML models | Gnina (CNN) | Grid of atom type channels representing protein-ligand complex in 3D space. | *McNutt et al. (2025); Francoeur et al. (2020)* |

We chose to exclude physics-based methods, such as free energy calculations, due to their prohibitive computational cost for large-scale screening, as per condition 4. Additionally, we limited our selection to tools that are open-source and installable via conda-forge, with the exception of Gnina, which is accessible through a Docker image.

### The coronavirus main protease binding pocket is conserved among viral homologs

The coronavirus main protease (M^pro^), also known as the 3C-like protase (3CLpro), M^pro^ is a highly conserved cysteine-protease and an attractive target for broad-spectrum antiviral development (***von Delft et al., 2023***). M^pro^ has a well-characterized mechanism of action, catalyzing eleven different site-specific cleavages within the two polyproteins translated from the first open reading frame (ORF1a/b) of the viral genomic RNA (***Dai et al., 2020***), a crucial step in the viral replication process. As such, M^pro^ was a primary focus during the early stages of the COVID-19 pandemic, with over 1,500 crystal structures now available in the protein data bank (***Jin et al., 2020***; ***Boby et al., 2023***). Furthermore, M^pro^ has no human homologs, and its substrate specificity is different from that of human cysteine proteases, reducing the risk of off-target effects and making it a promising target for successful clinical development. (***Dai et al., 2020***; ***von Delft et al., 2023***)

Being a key component of viral replication, M^pro^ is highly conserved across the coronavirus family. This conservation is evident in the MSA of all the seven coronavirus M^pro^s that infect humans, shown in Figure 2a. Exact residue matches (in red), and by aminoacid-type (in yellow, up to one missmatch, indicated in red font) are significant (49% of the reference sequence length), especially for residues in the binding pocket (indicated with blue boxes), even when overall sequence identity with the reference SARS-CoV-2 is as low as 41%. This sequence conservation is mirrored at the structural level. Mpro is a homodimer with three domains per monomer: I, II and II. The active sites are found at each interface of domains I and II and consists of four substrate-binding subpockets, S1’, S1, S2 and S4 Among these, S1, S2, and S4 are highly conserved across the viral family, as these support the catalytic core, composed of a cysteine-histidine dyad (***Yang et al., 2005***). Structural alignment of the folded models for SARS-CoV-1, MERS-CoV and HCoV-229E, with respect to SARS-CoV-2 in complex with Ensitrelvir (***Noske et al., 2023***) (Figure 2b), further highlights this structural conservation. Even for the most distantly related homolog to SARS-CoV-2 (pocket surface in white), HCoV-229E (pocket surface purple), the pocket structure remains largely unchanged, except for minor differences in S2.

This degree of structural conservation enables a ligand-transfer strategy, in which the ligand of a SARS-CoV-2 reference can be docked into the *apo* folded models of other homologs, for which no experimental crystal structure is available. More importantly, the structural conservation of the ligand pocket is a key condition for the development of broad spectrum antivirals. By preserving key protein-ligand interactions that are known to be important in the inhibition of SARS-CoV-2 (a target that has already been well-characterized), we can prioritize compounds likely to retain potency across diverse coronavirus targets, as well as guide the design of new molecules that would keep those interactions.

### The broad-spectrum pipeline is retrospectively validated with SARS-CoV-2 and MERS-CoV M^pro^

Using data generated by the ASAP Discovery Consortium, we retrospectively validated our predictive pipeline through a cross-target approach. The availability of X-ray crystal structures for both SARS-CoV-2 and MERS-CoV M^pro^ allowed us to use one as a template and the other for testing, enabling direct comparison between predicted and experimental P-L complexes on a per-compound basis. This validation process is illustrated in Figure 3a and was applied to a set of 43 compounds (the *validation set*) for which reference structures were available for both targets. As an example, we begin by selecting a SARS-CoV-2 M^pro^ P–L complex as a reference (here we show *ASAP-0008314*). Using this reference, we generate a folded and docked model of the MERS-CoV homolog. Since a MERS-CoV crystal structure with the same ligand is also available, we can compare the predicted model with experimental crystallographic data as a way to evaluate the performance of our model. In this case, the predicted protein structure aligns closely with the crystal structure (see insert in Figure 3b), though minor deviations are observed in the ligand pose. Conversely, we can also start with a MERS-CoV M^pro^ crystal template, to generate a model for the SARS-CoV-2 homolog, and compare it with the corresponding SARS-CoV-2 crystal structure. Following this pathway with *ASAP-0008314* leads to small differences in the binding pocket, but the ligand pose is well conserved (Figure 3c). By evaluating both prediction pathways, we can assess the overall reliability of the structural predictions that we can expect when the pipeline is applied to other viral homologs.

**Figure 3.**
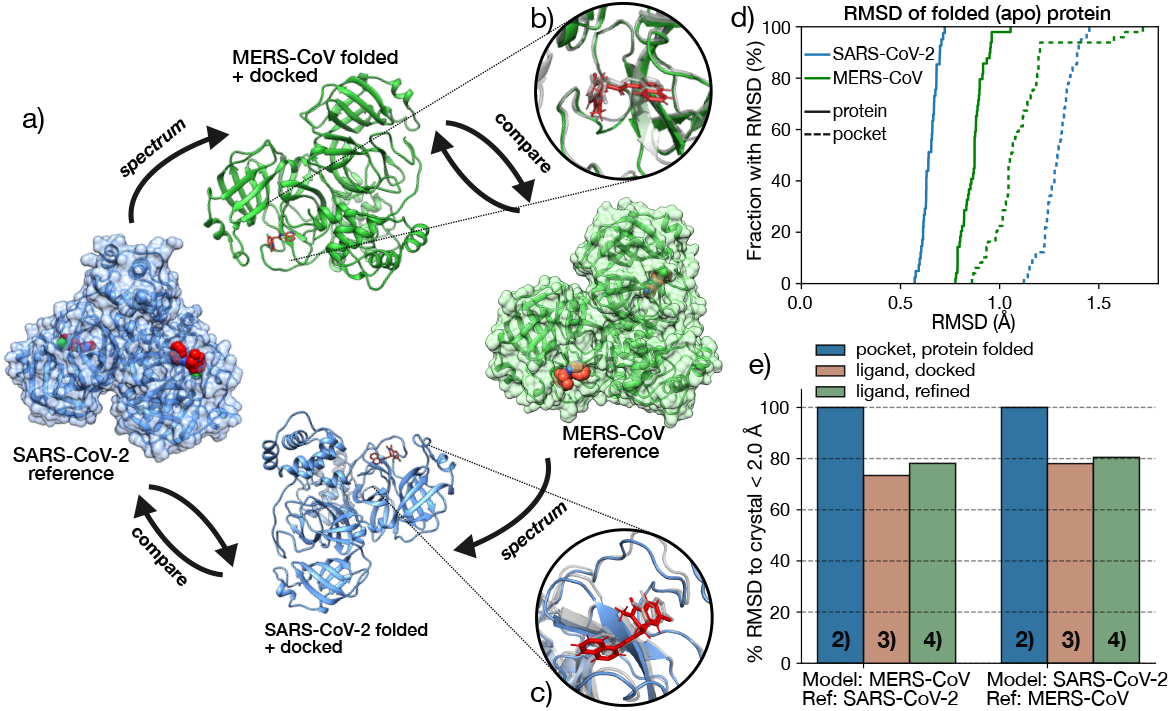
Our pipeline was retrospectively validated against a set of overlapping SARS-CoV-2 and MERS-CoV P-L co-crystalized structures. (a) Schematic illustration of the validation process. A reference SARS-CoV-2 M^pro^ P-L complex serves as a template for folding and docking a MERS-CoV apo-protein; the docked complexes are then compared with the X-ray crystal structures available for SARS-CoV-2, in complex with the same set of ligands. The same process can be followed to predict the SARS-CoV-2 structure from a MERS-CoV template. (b) A zoomed view of the binding pocket of the predicted MERS-CoV M^pro^ structure (green) compared with the SARS-CoV-2 template (shown here in complex with ASAP-0008314, gray). (c) Predicted SARS-CoV-2 (blue) starting from a MERS-CoV M^pro^ template (shown here in complex with ASAP-0008314, gray). (d) The ECDF cumulative distribution of the heavy-atom RMSD between the SARS-CoV-2 (blue) and MERS-CoV M^pro^ (green) apo-proteins predicted via AlphaFold2 and the corresponding crystal structure templates, shows good agreement between the two. The protein RMSD is shown in solid lines while the binding pocket (residues within <5.0Å from the ligand) RMSD is shown in dashed lines. (e) Fraction (in %) of RMSD within 2A with respect to the crystal structure with matching ligand for: the binding pocket of the AlphaFold folded proteins (blue), the ligand after the transfer docking (green) and the ligand after docking and refinement (red), corresponding to the 4, 5 and 6th steps of the pipeline in Figure 1, respectively. Here, we define the binding pocket as all residues within 5Åfrom the ligand.

To quantify structural accuracy, we first evaluate the protein folding step. Figure 3d shows the empirical cumulative distribution function (ECDF) of the RMSD values for the folded structures compared to the corresponding X-ray crystal structures, for the full protein (solid lines) and the binding pocket (dashed lines). The RMSD for the full protein is low and consistent across the samples for both targets, though slightly higher for the MERS-CoV predictions. In contrast, the binding pocket RMSDs are larger due to the smaller residue subset, particularly when predicting SARS-CoV-2 from a MERS-CoV template. This is evident from the example shown in the insert, where the binding pocket loops exhibit minor but notable deviations. Still, all RMSD values remain under 2.0Å, indicating reliable structural predictions.

We next validate and compare the accuracy of steps 2 to 4 of the pipeline—folding, ligand pose transfer, and MD refinement—by calculating the RMSD of the ligand pose with respect to the crystal reference. Figure 3e reports the percentage of structures with RMSD < 2Å for the predicted pocket (blue), transferred and docked ligand (green), and ligand after MD refinement (red). These are calculated separately for the two prediction pathways: SARS-CoV-2 to MERS-CoV (left), and MERS-CoV to SARS-CoV-2 (right). As discussed in Figure 3d, the folding step performs exceptionally well, with 100% of predicted pockets within 2Å of the reference. The accuracy drops for ligand pose prediction to 70–80%, but improves slightly after MD refinement, highlighting the need for energy minimization to resolve minor clashes introduced by pocket differences. Interestingly, the predictions are more accurate when starting from a MERS-CoV template (predicting a SARS-CoV-2 model), whereas the folding step had shown slightly better results in the reverse direction. This suggests that the discrepancy likely stems from the docking stage, and possibly from the POSIT algorithm itself, which may be biased toward targets more represented in its training data. Since POSIT uses both proprietary and public examples to predict pose quality, its performance could reflect underlying data availability or similarity.

Overall, the retrospective validation results demonstrate that our pipeline produces reliable protein and ligand structures, with the majority of predicted complexes closely matching the corresponding crystal structure. These predictions are expected to perform best in cases where the ligand adopts a similar binding mode across homologous targets.

### Protein-Ligand interaction fingerprints are recovered on the predicted models

To develop compounds that maintain potency across multiple viral homologs we must understand which P-L interactions are essential for inhibition, and conserve these interactions across the viral panel. During a reversible inhibition process, when a ligand binds to the binding pocket of a host protein, it is stabilized by different—non-covalent—intermolecular interactions. Those interactions can sometimes mimic the interactions between the protein and its substrate, and/or block the function of key residues on the active site, inhibiting the function of the protein as a result. Therefore, capturing patterns in those non-covalent protein ligand interactions is essential to understand the function of a protein, refining molecular docking or virtual screening poses and designing compounds that can inhibit the protein function (***Adasme et al., 2021***). Here, we attempt to do the latter, by establishing a relation between the interactions of a ligand with the binding pocket of a viral homolog protein target and its potency. Previous drug repurposing studies using PLIF-based virtual screening have shown that conserved protein–ligand interaction patterns can be linked to antiviral activity against SARS-CoV-2 (***Elfiky, 2020***; ***Schake et al., 2023***), suggesting that such interaction motifs can be used to prioritize repurposed compounds and scaffolds.

Starting from a reference P-L complex with known potency, it is useful to compare the PLIFs of that reference with those from folded homologs and docked models. If the ligand is known to be active against the reference target, then targets inhibited by that same ligand are likely to preserve key interactions critical for inhibition. Therefore, a ligand with broad spectrum activity against multiple targets should maintain a similar pattern of interactions with the conserved regions of the binding pocket across viral homologs. Figure 4a shows a 3D representation generated with Maestro (***Schrödinger, LLC, 2025***) of the binding pocket of SARS-CoV-2 M^pro^ with one of the ligands in the pipeline’s validation set, *ASAP-0008314*, for the crystal reference (left) and the predicted MERS-CoV model (right). We can see that for this example the pose is conserved in the two targets and, while the program only detected one hydrogen bond interaction in the SARS-CoV-2 pocket, namely, with Glu 166, the same Glu interaction is conserved in the MERS-CoV predicted model. Based on the interaction pattern, we could expect the compound to be active against both targets. Indeed, for *ASAP-0008314* a mean pIC_50_ of 6.70 ± 0.0659 was measured for SARS-CoV-2 M^pro^ and 5.69 for MERS-CoV. The Glu 166 hydrogen bond interaction is also conserved when we predict a SARS-CoV-2 model from a reference MERS-CoV M^pro^ crystal (Figure 4b). It is also interesting to compare the two prediction pathways. When we start from the MERS-CoV M^pro^ reference crystal (Figure 4b, left), the Glu 166 hydrogen bond interaction is conserved in the predicted SARS-CoV-2 model (right), but more interactions are detected by the visualization program. This observation partly explains the lower predicted pose accuracy observed Figure 3 b, as the poses seem to be slightly shifted between the SARS-CoV-2 reference and model.

**Figure 4.**
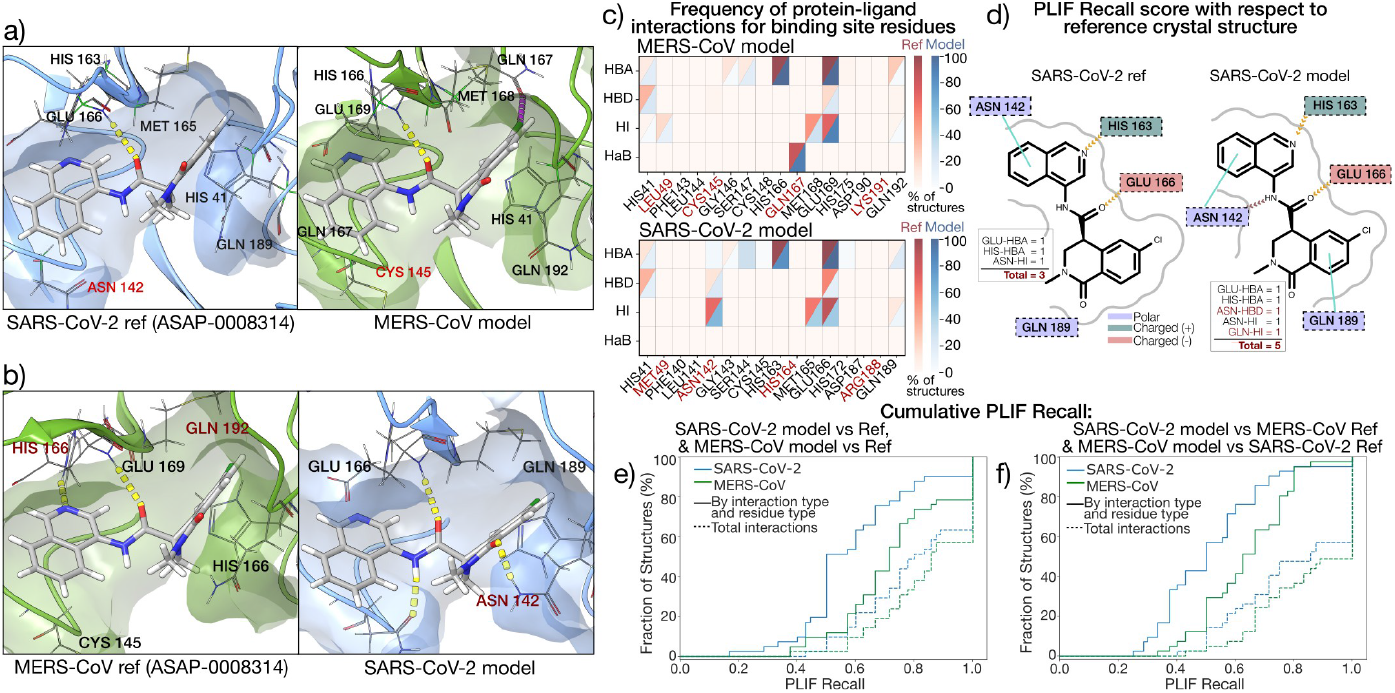
Protein-ligand interaction fingerprints are relatively conserved in the folded SARS-CoV-2 and MERS-CoV structures compared to the crystal. 3D visualization generated with Maestro of P-L hydrogen-bond interactions (yellow) of crystal structure vs docked pose for (a) MERS-CoV M^pro^ as predicted from a SARS-CoV-2 reference crystal, and (b) SARS-CoV-2 M^pro^ as predicted from a MERS-CoV reference, both in complex with *ASAP-0008314*. The 3D visualization in the top panels shows that key interactions are conserved as we go from SARS-CoV-2 to MERS-CoV, with some differences (red) as we go from MERS-CoV to SARS-CoV-2. (c) PLIFs for matching residues in the binding pocket of SARS-CoV-2 (top) and MERS-CoV (bottom), with unmatched residues indicated in red font. The frequency (in %) per residue and interaction type is shown as a red gradient for the set of X-ray crystal structures, and blue for the predicted models. Interaction types are Hydrogen Bond Acceptor (HBA), Hydrogen Bond Donor (HBD), Hydrophobic Interaction (HI) and Halogen Bond (HaB). Results show agreement between model and reference, and conservation of key interactions between SARS and MERS. (d) 2D visualization of PLIFs for the SARS-CoV-2 reference and predicted model, showing HBA, HBD and HIs in orange, brown and cyan, respectively, with residues also by side chain properties. The insert illustrates how interactions are accounted for in terms of *interaction type and residue type* and *total number of interactions*. (e) Cumulative distribution function (CDF) plot for the PLIF Recall score for ref vs model calculated as described in the text, for SARS-CoV-2 (blue) and MERS-CoV (green), and the two types of interaction match criteria illustrated in d). MERS-CoV models shows a better agreement with experiment than SARS-CoV-2, in terms of PLIFs.

The interaction pattern with the binding pocket for all ligands in the pipeline validation set is presented as a heatmap in Figure 4c, for the MERS-CoV (top) and SARS-CoV-2 M^pro^ (bottom) structures. The distribution for the crystal references is shown in red, while that of the predicted models is shown in blue. Interactions are calculated using the software PLIP (***Salentin et al., 2015***; ***Adasme et al., 2021***), with hydrogen bond acceptor (HBA), hydrogen bond donor (HBD), hydrophobic interactions (HI) and halogen bond (HaB) chosen to represent the fingerprints. The binding pocket residues were defined as those within 5Å from the ligand, and chosen to match spatially for SARS-CoV-2 and MERS-CoV M^pro^, with non-matching residues in red font. We can see certain key interactions are consistent between the two targets in the validation set: a histidine HBA (His 163 in SARS-CoV-2), a HI with a methionine (Met 165), and a HBA, HBD and HI with glutamine (Glu 166) that are also evidenced in panel a) and b). At the same time, some interactions are not conserved due to residue changes, such as those with Asn 142 in SARS-CoV-2, and Gln 167 in MERS-CoV. Notably, except for one interaction in the SARS-CoV-2 predicted model (Ser 144), the fingerprints are consistent between the reference and the model, meaning the prediction is generally accurate, especially for MERS-CoV M^pro^.

We devised a scoring method that compares the PLIFs of a P-L model against a specified reference in an efficient manner, therefore allowing the screening of compounds in search of one that maintains non-covalent interactions associated with the protein function. We assume that we should penalize the loss of interactions, rather than reward new interactions, as the interactions seen in the reference are supported by biochemical and structural data, while the interactions seen in the model are unverified. We therefore use a version of the Tversky similarity measure which is similar to recall. Given two sets, A and B, the Tversky index is defined as:

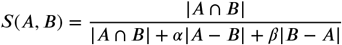

Setting *α* = 1 and *β* = 1 gives the symmetric Tanimoto coefficient (also known as the Jaccard index), while setting *α* = 1 and *β* = 0 gives:

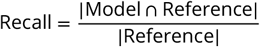

We then plot the empirical cumulative distribution function (ECDF) of the fraction of predicted models vs recall for each level of detail (Figure 4e,f). For instance, at the most general level (*Total interactions*), the recall is given by:

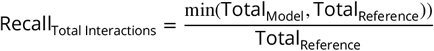

For the higher level of detail shown in Figure 4e-f (*By interaction type and residue type*), the intersection is calculated such that interactions are only counted the same if they are the same type (see Methods for more details) and involve the same amino acid in the protein.

We first calculated this recall score to compare the predicted models with their corresponding X-ray crystal structures (Figure 4e). In correspondence with the pocket RMSD values for SARS-CoV-2 and MERS-CoV (Figure 2), the recall scores for SARS-CoV-2 were lower than for MERS-CoV. This extends to the comparison of the predicted models to the references used to generate them (Figure 4), with the predicted models for SARS-Cov-2 retaining fewer of the interactions in the reference than the predicted models for MERS-CoV.

### Empirical and machine learning models are effective methods for scoring new ligands in SARS-CoV-2 and MERS-CoV M^pro^

Having validated the accuracy of ligand pose prediction in X-ray crystal structures shared between SARS-CoV-2 and MERS-CoV M^pro^, we next evaluate the binding affinity prediction capabilities of our pipeline. We focus on the scoring methods listed in Table 1, which include the empirical scor-ing functions ChemGauss4 (***OpenEye, Cadence Molecular Sciences, Inc., 2025***) and AutoDock Vina (***Eberhardt et al., 2021***), as well as the machine learning (ML) model Gnina (***McNutt et al., 2021***).

Empirical scoring functions are commonly used in molecular docking due to their efficiency, which makes them ideal for virtual screening campaigns. These models typically use simple descriptors to describe the interactions involved in P-L binding (e.g., steric and hydrogen bond interactions) and a regression or classification algorithm trained on experimental affinity datasets. For instance, ChemGauss4 is a purely Gaussian-based function, while AutoDock Vina is a hybrid scoring function, combining weighted Gaussian and linear terms. See Table 1 for details on their descriptors. In contrast, ML-based models such as Gnina are trained using non-linear functions capable of capturing more complex binding relationships. Like empirical models, they can have descriptors derived from structural features but often achieve improved generalization and predictive performance (***Liu and Wang, 2015***). For broad-spectrum antiviral screening, both scalability and generalizability are needed, making ML models promising scoring functions for this task. We compared the performance of these four scoring functions using experimental binding affinity data (pIC_50_) the set of overlapping SARS-CoV-2 and MERS-CoV M^pro^ P-L complexes, introduced in the previous section. Additionally, we ran an additional set of calculations for the P-L complexes for which we have a crystal structure of either SARS-CoV-2 or MERS-CoV, but not necessarily a corresponding structure for the other target. This amounted to a total of 82 and 162 unique crystal templates of SARS-CoV-2 or MERS-CoV, respectively. After docking and filtering, we retained 78 and 149 compounds for SARS-CoV-2 and MERS-CoV, respectively. Pearson and Kendall’s *τ* correlation coefficients between predicted and experimental pIC_50_ values are shown in Figure 5a, with confidence intervals estimated via bootstrap resampling indicated for each case. Overall, Gnina consistently outperforms other methods in both targets, while ChemGauss4 shows weak correlation. However, we note that low correlation across methods may stem from differences in scaling between the different scoring methods, as well as the small dataset size—also evident in the large error bars.

**Figure 5.**
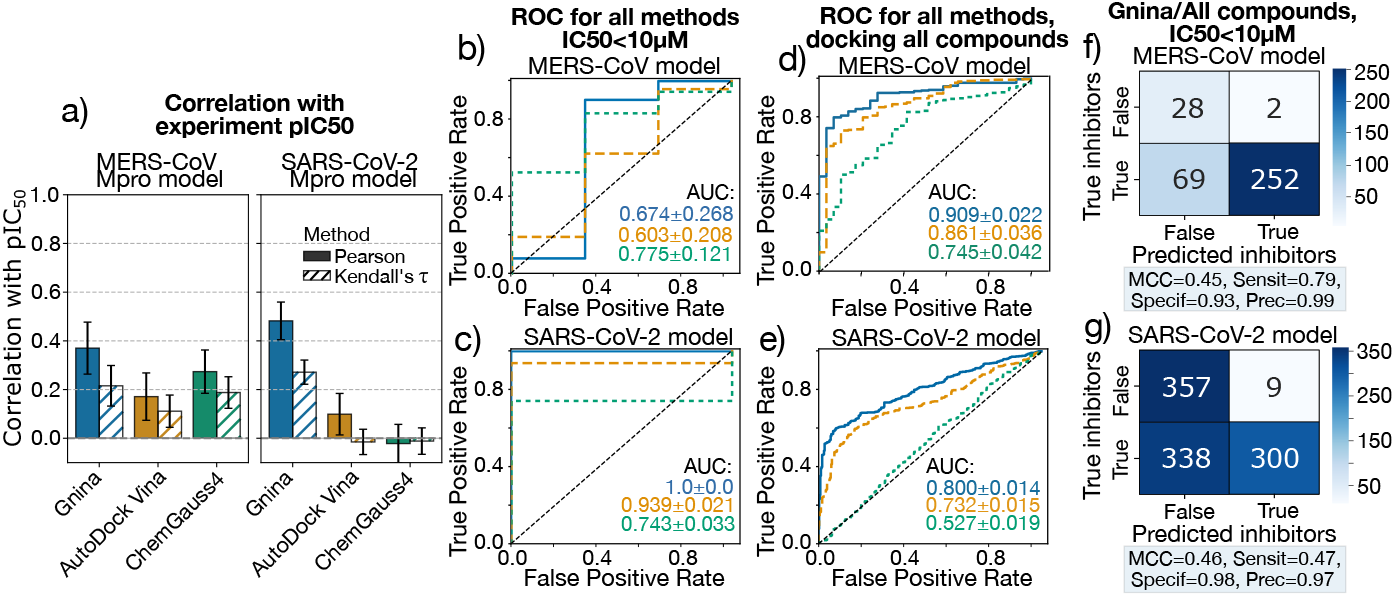
Calculated Scores for the folded MERS-CoV and SARS-CoV-2 M^pro^ targets show a direct correlation with antiviral efficacy as determined from biochemical assays. (a) Pearson (solid bars) and Kendall’s *τ* (hatched bars) correlations between experimental pIC_50_ values for MERS-CoV and SARS-CoV-2 M^pro^ inhibitors, and scores predicted for the folded and docked models (78 and 149, respectively). Error bars show 95% confidence intervals from bootstrap resampling. (b-c) Receiver operating characteristic (ROC) curves for classification in the same ligand set using Gnina (blue), AutoDock Vina (orange) and ChemGauss4 (green), for the MERS-CoV (b) and SARS-CoV-2 (c) predicted models. AUC scores are reported, with errors calculated via bootstrap resampling. Here, a ligand is classified as an “inhibitor” when the IC_50_ is below 10*μ*M. (d–e) ROC curves for the extended set predicted by docking all the ligands in the *ASAP-0008314* folded model (351 and 1004). (f-g) Confusion matrix for the Gnina CNN predicted affinities for the MERS-Cov (f) and SARS-CoV-2 (g) models, and all available compounds docked. The Matthews Correlation Coefficient (MCC) score, sensitivity (Sensit), specificity (Specif) and precision (Prec) are shown in a shaded box below each matrix.

Because active vs inactive classification is often more relevant than absolute prediction in early drug discovery, we also assessed model performance using receiver operating characteristic (ROC) curves, which assess how well the model distinguishes inhibitors across varying thresholds (Figure 5b–c). Ligands with IC_50_ < 10*μ*M were labeled as “inhibitors”. The area under the ROC curve (AUC) provides a threshold-independent measure of classification performance, which can be used to assess the model’s ability to prioritize active compounds. Notably, Gnina achieves perfect classification (AUC = 1) for the SARS-CoV-2 model folded from a MERS-CoV reference, while AutoDock Vina achieves similar performance at lower computational cost, which could provide advantageous in large virtual screening campaigns. In general, the AUC score seems to capture the performance of the evaluated scoring methods more accurately than the linear correlation, specially if these methods are to be applied for early-stage prioritization. Note that the ROC curve for the SARS-CoV-2 model is based on a smaller dataset (78 compounds vs. 149 for MERS-CoV), which may give the curve a more “rectangular” appearance due to fewer points. However, the bootstrap confidence intervals suggest that the AUC values remain robust despite the reduced sample size.

We then expanded the test set to include all compounds with experimental pIC_50_ values, regardless of crystal structure availability. Using a single folded model, namely, that for *ASAP-0008314*, we docked and refined 351 and 1004 compounds for MERS-CoV and SARS-CoV-2, respectively (Figure 5d–e). ROC curves from this extended set also show strong predictive power for Gnina (AUC =0.90 and 0.80). Confusion matrices for Gnina predictions (Figure 5f–g) indicate high specificity and precision across both targets, meaning the model can correctly identify most of the true inactives correctly. Interestingly, performance trends reversed in the extended set: the MERS-CoV model outperformed SARS-CoV-2 across-the-board. This may reflect differences in ligand pose accuracy (RMSD of 0.5Å vs. 1.8Å, respectively) or biases in training data. Specifically, both Vina and Gnina were trained using iteration of PDBbind prior to 2020, before an influx of SARS-CoV-2 structures from the COVID Moonshot project (***Boby et al., 2023***).

In summary, both Gnina and AutoDock Vina demonstrate strong potential for potency prediction in our sequence-to-structure pipeline, showing utility across transfer-docked and *de novo* docked complexes. Their predictive accuracy and computational efficiency make them viable tools for structure-based discovery of broad-spectrum antivirals.

### The spectrum breadth of affinity prediction is evaluated across a diverse coronavirus panel

To assess the broad-spectrum potential of our predictive pipeline, we evaluated its performance on a diverse panel of coronaviruses beyond SARS-CoV-2 and MERS-CoV. Our analysis builds on the experimental results reported by ***Leonard et al***., who developed a sensitive and low-background fluorescent assay based on a FlipGFP variant. This assay enables the measurement of protease activity for a wide range of viral M^pro^ targets and was applied to a panel of both human and zoonotic coronaviruses spanning the alpha, beta, delta, and gamma genera. Evaluating our pipeline across this expanded coronavirus panel, which not only includes all known human-infecting strains (Figure 2), but also animal viruses with zoonotic potential, is crucial for designing preventive therapeutics with some level of efficacy against viral homologs that may emerge in the future.

To that end, we initialized our pipeline using a SARS-CoV-2 crystal structure of M^pro^ bound to Ensitrelvir (*PDB: 8dz0* (***Noske et al., 2023***)) and applied it to the 16 M^pro^ targets depicted in the phylogenetic tree in Figure 6a. Note that although SARS-CoV-2 is the reference structure for ligand transfer, it is also included in the evaluation panel. To maintain consistency, the SARS-CoV-2 protein sequence output from the BLAST search is folded and docked with Ensitrelvir following the same pipeline as the other 15 coronaviruses, ensuring all targets are evaluated under the same criteria. Additionally, we excluded gamma-Coronaviruses from this study, due to their lower zoonotic risk, which also allowed for a more streamlined analysis. The experimental fluorescence signal was converted into a “signal inhibition” estimate by subtracting 100%−[normalized reporter signal],yielding a value directly proportional to expected binding affinity. Figure 6b presents the Pearson and Kendall’s *τ* correlations between the experimental signal inhibition and predictions from each scoring method listed in Table1. Confidence intervals from bootstrap resampling are shown for each case. For most methods, Pearson correlations are higher than Kendall’s *τ*, which does not exceed 0.5 for any metric. This suggests that while some methods exhibit moderate linear agreement with experimental data, they may not preserve the correct ranking of compounds across the panel. We also see that the strongest correlations (both Pearson and Kendall’s) were observed for sequence similarity scores, for both the binding pocket and the full protein, likely due to high potency of Ensitrelvir against SARS-CoV-2 (i.e., nearly complete signal inhibition). Similarly, the protein–ligand interaction fingerprint (PLIF) scores also showed high Pearson correlation, while ligand RMSD performed poorly, indicating that interaction-based and sequence-based features are more predictive of inhibition in this context than just ligand posing. Among the empirical and machine learning scoring functions, Gnina and AutoDock Vina exhibited modest correlations with experiment, consistent with our previous observations in the SARS-CoV-2/MERS-CoV validation set. These low correlations may arise from a difference in scaling of the outputs and model generalization that cannot be captured by linear analysis alone. To better evaluate classifier performance, we plotted signal inhibition (log-transformed) against Gnina and PLIF scores in Figure 6c, and show their corresponding confusion matrices in panel d. The experimental threshold for inhibition was defined as >50% of signal inhibition. Model predictions, however, have different scaling and therefore it is hard to pick a single cutoff that would work with every estimate of ligand potency. Therefore, we selected cutoffs to evaluate ability of each method to avoid false negatives while testing for false positives (i.e., testing for specificity while setting sensitivity to 1.0): 7.2 kcal/mol for Gnina and 0.64 for PLIF, indicated by red dashed lines. Experimentally, only three targets, namely, SARS-CoV-2, SARS-CoV-1, and MERS-CoV, show significant inhibition by Ensitrelvir. The PLIF recall accurately identified all three without false positives, achieving perfect classification (MCC = 1.0). In contrast, Gnina misclassified four additional targets as inhibited, including HKU1 (which does show partial inhibition at ∼ 33%). As a result, Gnina had lower specificity (0.69) and precision (0.43). While PLIF yielded ideal classification, it did not preserve the correct ranking of the compounds, explaining its lower Kendall’s *τ* coefficient. This contrast between a low Kendall’s *τ* correlation and high classification performance indicates that scoring models such as PLIF can correctly classify active compounds while still misranking them within the active and inactive groups. This suggests that, despite limited ranking precision, such models remain effective for distinguishing actives from inactives, which is a useful feature for early-stage hit prioritization. Figure 6e presents the ROC curves and AUC scores for all scoring methods, with color schemes matching panel a. As expected, the AUC trends mirror the previous trends but offer a more robust metric for evaluating classification of active vs inactive targets. In this case, we see that both PLIF and the binding pocket sequence similarity scores are perfect classifiers with an AUC of 1.0, which highlights the strong correlation between sequence and structure in the selected dataset. On the other hand, neither empirical and ML methods can fully capture this relationship, and incorrectly predict some targets to be inhibited by Ensitrelvir. Still, both Gnina and AutoDock Vina showed strong classification performance, suggesting that with appropriate calibration, they could be prospectively applied to prioritize new compounds across coronaviruses. Note that the evaluated set remains relatively small, as reflected by the larger uncertainties in the predicted outputs. More conclusive insights into broad-spectrum performance will require extending this analysis to additional compounds and experimental data. To further assess the pipeline’s potential for prospective prediction, we extended the analysis to 43 ASAP Discovery compounds, specifically, those with available X-ray co-crystal structures in both SARS-CoV-2 and MERS-CoV. For each compound, we generated protein–ligand models for all 16 targets and calculated PLIF, Gnina, and AutoDock Vina scores. The score distributions are shown in Figure 6d, with the total number of compounds that were successfully docked and refined indicated for each target. Notably, PLIF scores displayed the largest variability across targets, as evidenced by wide interquartile ranges and large error bars. This inconsistency may be due to differences in interaction with the compounds across diverse targets, as well as varying potency for the reference, SARS-CoV-2, for different compounds. This observation may indicate that reference-based scoring methods may not be as effective when assessing compounds that could have broad-spectrum potency, but reduced activity against the reference target. In contrast, Gnina and AutoDock Vina scores were more consistent, reinforcing the value of empirical and ML for ranking diverse targets. Based on these metrics, a practical strategy for identifying broad-spectrum candidates is to prioritize compounds with consistent high predicted potency across the panel (particularly those with scores comparable to SARS-CoV-2) for further experimental validation.

**Figure 6.**
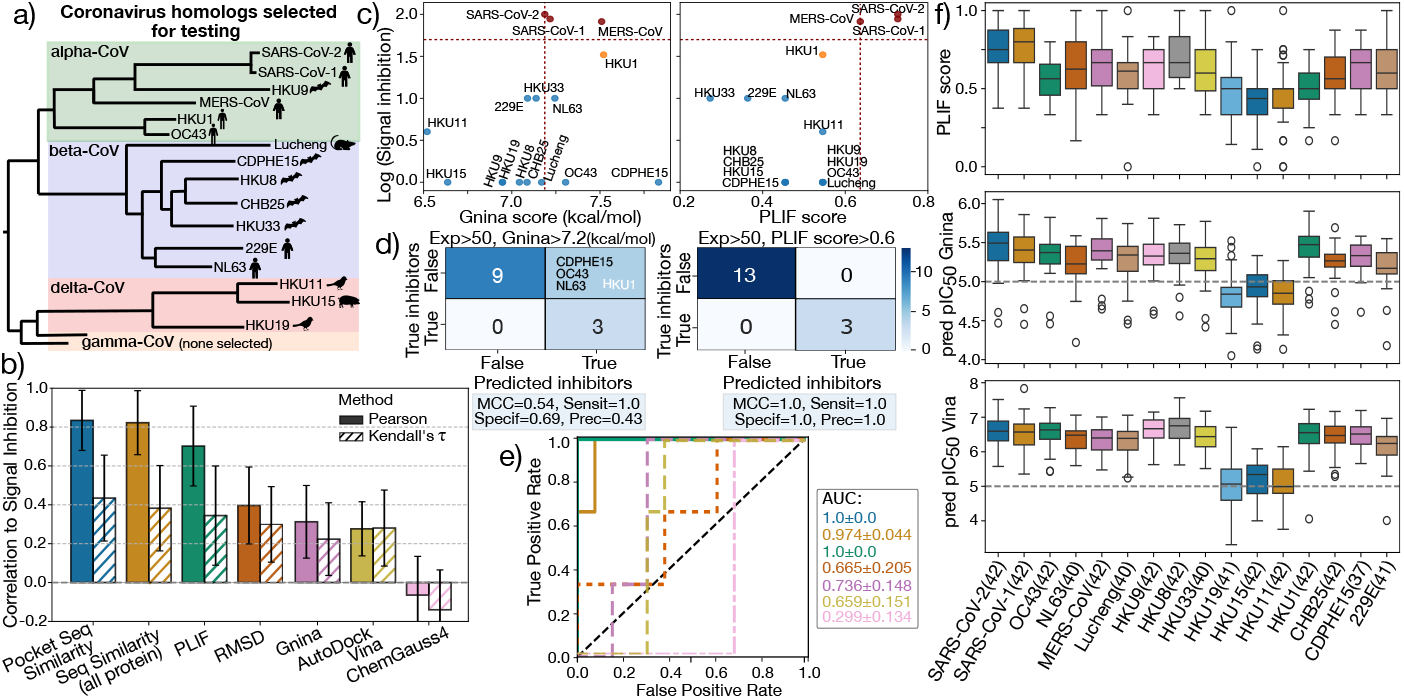
Broad-spectrum activity is tested against 16 human and non-human coronaviruses, showing accurate prediction of signal inhibition in Ensitrelvir fluorescence-based assay. (a) Phylogenetic tree of alpha, beta and delta coronaviruses tested against our pipeline, adapted from Fig 4 in (***Leonard et al., 2023***).(b) Pearson (solid bars) and Kendall *τ* (hatched bars) correlations between recovered signal (defined as 100%-[Normalized reporter signal], as presented in cell-based experimental assay) and binding affinity predictions from the different scoring methods studied in this manuscript: Binding pocket and all-protein sequence similarity to SARS-CoV-2 reference, PLIFs (by interaction type and residue type), Ligand RMSD with respect to the reference crystal, Gnina CNN score, AutoDock Vina and ChemGauss4 score. c) Scatter plot of reporter signal inhibition (in the log10 scale) vs Gnina (left) and PLIF (right) predicted score for all 16 targets. Dashed red lines indicate the cutoff set for labeling compound activity, while inhibited, partially and non-inhibited compounds, according to experiment, are indicated with red, orange and blue, respectively. d) Confusion matrix for Gnina (left) and PLIF (right) score predictions, with predicted inhibition cuttoff of 7.2 kcal/mol, and 0.6, respectively, and three true active (red in panel b). e) ROC curves for each of the scoring functions and AUC scores, with colors matching those in panel a). f) Distribution of predicted PLIF scores (top), Gnina pIC_50_s (middle) and AutoDock Vina pIC_50_s (bottom), across 16 CoV targets for the set of 43 ASAP compounds with available SARS-CoV-2 and MERS-CoV X-ray crystal structures. Outliers are indicated as unfilled circles.

However, we also observed that the delta-CoV targets, HKU19, HKU11 and HKU15, where repeatedly scored lower for the compound set. While we have yet to confirm this behavior with additional experiments on these molecules, it could be an indication that our method cannot generalize to targets that are less evolutionary related to SARS-CoV-2. A potential fix could be to retrain the Gnina ML model to identify specific patterns of that genera. Another possibility is that the structure prediction of the deltacoronaviruses is not accurate on the binding site, possibly due to the use of SARS-CoV-2 as a template. However, the binding domains of the crystal structure of HKU15 and alpha-CoVs should be well-conserved, with an RMSD<2Å (***Wang et al., 2022***).

In practice, these prospective predictions can be used to prioritize compounds for further experimental testing, particularly those scoring highly across multiple targets (using robust metrics like Gnina and AutoDock Vina). Subsequently, structure–activity relationship (SAR) could be used to refine lead compounds with similar interaction patterns, ultimately aiding the development of broad-spectrum antivirals.

We also evaluated how our ligand transfer and minimization strategy performs against state-of-the-art protein–ligand structure prediction approaches, specifically the co-folding methods Chai-1 and Boltz-2. As shown in Figure 7, our models achieve comparable ROC AUC values across five scoring metrics, demonstrating similar classification performance across the coronavirus panel. In particular, our method performs particularly well with structurally driven scores such as PLIF and ligand RMSD, and the ML-based Gnina model, while co-folding methods show stronger results for empirical scoring functions like AutoDock Vina and ChemGauss4. The Boltz-2 model also includes a built-in affinity predictor, which achieved an AUC of 0.799±0.142, although this scoring function is specific to that software and cannot be applied to score other models. For robust prospective predictions, we recommend using an ensemble of models and scoring strategies, prioritizing compounds that exhibit consistent broad-spectrum activity across different structure-prediction and scoring methods.

**Figure 7.**
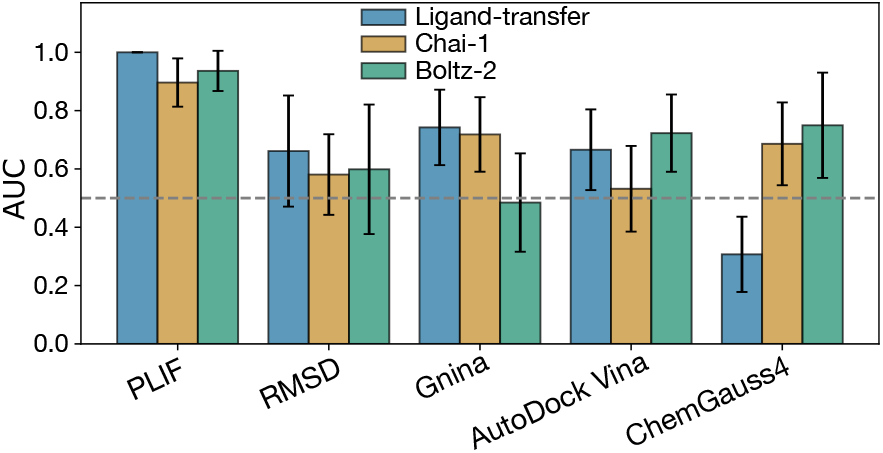
The ligand transfer and minimization strategy has comparable performance to state-of-the-art co-folding methods. Area under the ROC curve (AUC) for selected scoring methods applied to protein–ligand models of Ensitrelvir across the coronavirus panel. Results are shown for models generated using our ligand transfer and refinement approach (blue), and compared with two co-folding–based methods: Chai-1 (yellow) and Boltz-2 (green). The baseline of AUC=0.5, which corresponds to random classification performance, is shown as a gray dashed line.

## Discussion

The COVID-19 pandemic underscored the urgent need for novel antiviral strategies that can be rapidly deployed against emerging pathogens. At the beginning of the pandemic, research largely centered on re-purposing existing therapeutics as a quick and inexpensive strategy to deal with the sudden outbreak. However, these efforts were largely unsuccessful (***Edwards and Hartung, 2021***; ***Tummino et al., 2021***), in part because most molecular libraries are optimized for host-targeting compounds. More fundamentally, re-purposing strategies that ignore structural context and PLIFs fall short, as different drugs act through distinct mechanisms that cannot be generalized for every host and every virus.

Instead, pairing the knowledge of cross-target active compounds with structural insights can enable a more effective and rational drug discovery process, especially if a sudden outbreak occurs. Our structure- and interaction-informed pipeline would allow for re-purposing compounds known to be active for SARS-CoV-2 and testing prospectively against other coronavirus homologs, with scoring methods that combine data-driven empirical and ML techniques with structure-based interaction metrics. Beyond re-purposing, this pipeline can also be applied to designing molecules de-novo or refining existing scaffolds, leveraging structural knowledge to propose candidates for large-scale in silico screening. Paired with downstream refinement through more accurate, albeit expensive, computational methods and experimental validation, our workflow can become a valuable tool for hit identification (by screening large data sets against a panel of viral homologs), or even for early-stage lead optimization (by facilitating potential compounds or scaffold refinement based on cross-target efficacy).

Our results demonstrate that a structure-based pipeline integrating folding, docking and scoring can reliably predict the potency of protein–ligand complexes across diverse panel of coronavirus targets. In the retrospective validation against SARS-CoV-2 and MERS-CoV M^pro^, the pipeline accurately reproduced ligand poses and preserved key protein-ligand interaction features, as evidenced by low pose RMSD and consistent PLIF patterns. In this context, where sequence homology between targets is high, scoring approaches based on sequence and interaction similarity performed particularly well. Additionally, ML- and empirical-based scoring methods such as Gnina and AutoDock Vina showed strong predictive power, likely due to the close match between these targets and the training data used in the models. Similarly, in the broad-spectrum evaluation with Ensitrelvir, both reference similarity-based and data-driven metrics accurately identified which targets were inhibited (SARS-CoV-2, SARS-CoV-1, and MERS-CoV), likely because all three had strong sequence homology with the template, SARS-CoV-2. In contrast, scoring methods like ChemGauss4 did not successfully predict potency in any of the validation tests. While computationally efficient, such methods may lack sufficient accuracy for identifying potent inhibitors across a wide phylogenetic spectrum. Thus, we recommend a combination of efficient ML- or empirical scoring methods (e.g., AutoDock Vina, Gnina) with structure-informed approaches for fast and reliable prediction of broad-spectrum activity Our prospective evaluation with Ensitrelvir also highlights the strengths and limitations of different scoring strategies. Pose-based and PLIF metrics are particularly effective when the ligand is a strong inhibitor of the reference complex, as high-affinity interactions tend to be conserved across homologs, resulting in high classification accuracy (e.g., PLIF AUC = 1). However, these approaches may be less reliable when evaluating compounds with weaker binding to the reference, where structural conservation alone does not ensure cross-target activity. In such scenarios, empirical and ML methods are essential for generalization. We have yet to test our pipeline against such systems, due to the availability of experimental data. These limitations were also evident in our prospective application of the pipeline to other ASAP Discovery compounds, where predictions for more distantly related targets, particularly those from the delta-CoV genus, were less accurate, reflecting the challenges of generalizing beyond closely related homologs.

To address these limitations, we are actively expanding our validation efforts using new exper-imental data across a broader range of targets and compounds. In particular, we aim to evaluate prospective predictions from our structure-based pipeline in collaboration with experimental partners. Simultaneously, we are refining ML-based scoring models by incorporating protein–ligand interaction features explicitly and retraining them on coronavirus-specific datasets. This approach will enhance the ability of our pipeline to generalize to more phylogenetically diverse targets beyond the alpha- and beta-CoV genera. In parallel, we are exploring the integration of co-folding prediction models, which showed comparable classification performance to the ligand-transfer and refinement strategy presented here, while also offering improved computational efficiency— in terms of wall-time—by unifying the folding and docking steps. Furthermore, unlike ligand trans-fer–based methods, co-folding approaches do not require a conserved binding pocket, making them especially valuable for modeling targets with substantial sequence or structural divergence in the active site. Together, these findings and ongoing developments position our pipeline as a flexible and extensible framework for the structure-based prediction and prioritization of broadspectrum antiviral compounds, providing a valuable tool for future pandemic preparedness.

## Materials and Methods

### Sequence to structure modeling pipeline

In the first step of the piepline, the BLAST search was performed by querying NCBI’s QBLAST server(***Altschul et al., 1990***) to perform a sequence-alignment-based search to identify homologous protein sequences. The reference protein sequence was extracted from a PDB file containing the X-Ray Crystal structure for the protein-ligand complex, the BioPython. The BioPython PDB module (***Hamelryck and Manderick, 2003***) was used to extract the PDB sequence, and then the Blast module (***Cock et al., 2009***) was used to query the NCBI. After the homolog protein sequence are retrieved, the sequences are filtered by host information. Viral host and organism information is obtained from the NCBI Entrez database (***Schuler et al., 1996***; ***Sayers et al., 2021***), using the BioPython Entrez module. For section Referencessec:broad-ensit, the sequences are filtered based on the coronavirus panel in ***Leonard et al***..

After the filtering step, the protein sequences are folded using AlphaFold2 Multimer (***Evans et al., 2021***) with the reference protein as a template, as implemented in ColabFold(***Mirdita et al., 2022***). The coordinates of the reference ligand are transferred into the folded apo-proteins, and the protein-ligand complex is docked using the POSIT algorithm (***Kelley et al., 2015***), as implemented in the OpenEye Toolkit (***OpenEye, Cadence Molecular Sciences, Inc., 2025***; ***OpenEye, Cadence Molecular Sciences, 2024***). The docked poses are then refined by MD minimization, using OpenMM (***Eastman et al., 2023***). The docked and refined poses can then be employed for docking more compounds and scored using the methods described in the manuscript.

The co-folding models were prepared using the same protein sequences used for the AlphaFold2 step in our pipeline, as implemented in the Chai-1 (***Chai Discovery, 2024***) and Boltz (***Passaro et al., 2025***) software, with the co-folded structures then used as input for the scoring step. For the Chai-1 model, a SARS-CoV-2 crystal structure in complex with Ensitrelvir was provided as a template (***Noske et al., 2023***).

### Collection and Preparation of SARS-CoV-2 and MERS-CoV M^pro^ Experimental Data

Protein constructs for SARS-CoV-2 M^pro^ and MERS M^pro^ X-ray crystallography were expressed and purified at the Centre for Medicines Discovery, University of Oxford. X-ray crystal structures were determined via soaking or co-crystallisation at Diamond Light source according to the protocols outlined for SARS-CoV-2 M^pro^ (***Balcomb et al., 2025b***) and MERS-CoV M^pro^ (***Balcomb et al., 2025a***) respectively.

M^pro^ inhibition was measured using an in-vitro dose response assay performed at the Weizmann Institute of Science. A protocol involving a fluorescent M^pro^ substrate was used with slight differences in protocol employed between SARS-CoV-2 M^pro^ (***Lahav and Haim Barr, 2025b***) and MERS-CoV M^pro^ (***Lahav and Haim Barr, 2025a***). IC_50_s were determined by fitted to dose response curves, again with differing protocols. Both SARS-CoV-2 M^pro^ and MERS-CoV M^pro^ are homodimeric but differ in their dimerization kinetics. SARS-CoV-2 M^pro^ was fit to the Hill equation (***Hil, 1910***) using a four parameter logistic function. Due to MERS-CoV M^pro^’s weakly bound dimer, which can undergo ligand induced dimerization, an extension of the regular curve fitting procedure was requiring to mitigate spurious low-concentration activation.

Experimental inhibition data for the coronavirus panel in complex with Ensitrelvir were obtained from Leonard et al. (***Leonard et al., 2023***). The authors employed a FlipGFP fluorescence-based assay to quantify main protease activity in live cells. In this assay, fluorescence values were normalized with respect to a DMSO vehicle, such that decreased signal corresponds to higher protease inhibition, providing a quantitative measure of compound potency across viral targets. A value proportional to compound potency was obtained by subtracting the normalized fluorescence signal from 100% (the DMSO control)

### Calculation of protein-ligand interaction fingerprints

Protein-ligand interactions were calculated using the conda-installable PLIP software (***Salentin et al., 2015***; ***Adasme et al., 2021***). The code to convert the results of these interactions was influenced by PLIPify (***Lab, 2021***). The possible interaction types include Hydrogen Bond Donor, Hydrogen Bond Acceptor, Hydrophobic Interaction, Pi Stacking, Halogen Bond, and Salt Bridge.

As described in the main text, we assume that we should penalize the loss of interactions, rather than reward new interactions, as the interactions seen in the reference are supported by biochemical and structural data, while the interactions seen in the model are unverified. We therefore use a version of the Tversky similarity measure which is similar to recall. Given two sets, A and B, the Tversky index is defined as:

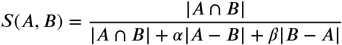

Setting *α* = 1 and *β* = 1 gives the symmetric Tanimoto coefficient (also known as the Jaccard index), while setting *α* = 1 and *β* = 0 gives:

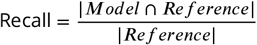

We then plot the empirical cumulative distribution function (ECDF) of the fraction of predicted models vs recall for each level of detail (Figure 4e,f). For instance, at the most general level (*Total interactions*), the recall is given by:

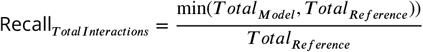

For the higher level of detail shown in Figure 4e-f (*By interaction type and residue type*), the intersection is calculated such that interactions are only counted the same if they are the same type and involve the same amino acid in the protein.

### Scoring Methods

To estimate the potency of ligand across the different viral homologs, we chose a total of six scoring functions, spanning three different types: Reference similarity-based functions (Sequence similarity of bonding pocket and whole protein, PLIF recall and ligand RMSD), Empirical-based (Chem-Gauss4 and AutoDock Vina), and ML-based scoring functions (Gnina). To estimate the sequence identity between a given protein with the reference, we divide the number of matching residues, by the length of the query protein:

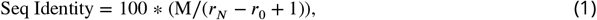

Where *M* is the number of matching residues, and *r*_*N*_ and *r*_0_ are the last and first residues of the query protein, respectively. To calculate the sequence similarity of the binding pocket, we first defined the binding pocket of the reference and query protein as all the residues within 4.5 Å from the ligand. Then, a pairwise alignment (as implemented in the BioPython pairwise2 module (***Cock et al., 2009***) is performed between the binding pocket residues of the query and the reference to find the number of matching residues. The equation 1 is applied to the residues in the binding pocket (i.e., *r*_*i*_ is indexed for the selected residues). The ligand RMSD is calculated using the OERMSD tool implemented in OpenEye Tools (***OpenEye, Cadence Molecular Sciences, 2024***), while the PLIF Recall score is described in detail on the main text. The ChemGauss4 score was calculated for the docked poses as implemented in OEDocking (***OpenEye, Cadence Molecular Sciences, Inc., 2025***), while AutoDock Vina was applied using the implementation in (***Trott and Olson, 2010***; ***Eberhardt et al., 2021***). For Gnina, the code as described in (***McNutt et al., 2021, 2025***) was employed, with the `crossdock_default2018`_[*i*]_ model used for the binding affinity prediction (***Francoeur et al., 2020***), where *i* corresponds to different seeds. Seeds 1-4 were applied to the ligand pose and averaged for a final affinity score.

## Data and Code Availability

The `drugforge` toolkit code is open-source and available on GitHub [https://github.com/choderalab/ drugforge] under a permissive MIT License. Additionally, a Jupyter notebook with instructions to run the sequence-to-structure pipeline, and scripts and data used to generate the plots are also available at https://github.com/choderalab/broad-spectrum-asap-paper, with data available under the CC0 License.

## Acknowledgments

The authors are grateful to the ASAP Discovery Consortium [http://asapdiscovery.org] and its numerous talented scientists for their contributions to data generation, scientific motivation, and many thoughtful discussions in the formulation of this work.

The authors thank Sukrit Singh, Benjamin Kaminow, Chris Iacovella, Mike Henry, Josh Horton and Karla Kirkegaard for helpful discussions during the preparation of this manuscript.

## Disclaimer

The content is solely the responsibility of the authors and does not necessarily represent the official views of the National Institutes of Health.

## Funding

Research reported here was supported in part by NIAID of the National Institutes of Health under award number U19AI171399.

JDC acknowledges support from NIH grant U19 AI171399, NIH grant P30 CA008748, and the Sloan Kettering Institute. MAC acknowledges support from the Sloan Kettering Institute and the Gordon MERIT Fellowship. AMP acknowledges support from NIH grant T32 GM115327.

## Disclosures

JDC is a current member of the Scientific Advisory Board of OpenEye Scientific Software. JDC has equity in and serves as the Chief Executive Officer of Achira, Inc., which is engaged in the creation of open foundation simulation models for drug discovery. The Chodera laboratory receives or has received funding from multiple sources, including the National Institutes of Health, the National Science Foundation, the Parker Institute for Cancer Immunotherapy, Relay Therapeutics, Entasis Therapeutics, Silicon Therapeutics, EMD Serono (Merck KGaA), AstraZeneca, Vir Biotechnology, Bayer, XtalPi, In3terline Therapeutics, the Molecular Sciences Software Institute, the Starr Cancer Consortium, the Open Force Field Consortium, Cycle for Survival, a Louis V. Gerstner Young Investigator Award, and the Sloan Kettering Institute. A complete funding history for the Chodera lab can be found at http://choderalab.org/funding.

## Author Contributions

Conceptualization: MAC, AMP, JS, JDC; Methodology: MAC, AMP, JDC; Software: MAC, AMP, HMO, IP; Formal analysis: MAC, AMP, JS, JDC; Investigation: MAC, AMP, JS, BHB, EJG, DF, HB, NL, DC, JS, RR, BL; Writing–Original Draft: MAC; Writing—review & editing: MAC, AMP, JS, HMO, IP, BHB, EJG, JDC; Funding Acquisition: JDC; Resources: JDC; Supervision: JDC.

## Notes

### Competing Interest Statement

John D Chodera is a current member of the Scientific Advisory Board of OpenEye Scientific Software. John D Chodera has equity in and serves as the Chief Executive Officer of Achira, Inc., which is engaged in the creation of open foundation simulation models for drug discovery. The Chodera laboratory receives or has received funding from multiple sources, including the National Institutes of Health, the National Science Foundation, the Parker Institute for Cancer Immunotherapy, Relay Therapeutics, Entasis Therapeutics, Silicon Therapeutics, EMD Serono (Merck KGaA), AstraZeneca, Vir Biotechnology, Bayer, XtalPi, In3terline Therapeutics, the Molecular Sciences Software Institute, the Starr Cancer Consortium, the Open Force Field Consortium, Cycle for Survival, a Louis V. Gerstner Young Investigator Award, and the Sloan Kettering Institute.

## References

PROCEEDINGS OF THE PHYSIOLOGICAL SOCIETY: January 22, 1910. The Journal of Physiology. 1910; 40(Suppl):i–vii. https://physoc.onlinelibrary.wiley.com/doi/abs/10.1113/jphysiol.1910.sp001386, doi: 10.1113/jphysiol.1910.sp001386.

Abramson J, Adler J, Dunger J, Evans R, Green T, Pritzel A, Ronneberger O, Willmore L, Ballard AJ, Bambrick J, et al. Accurate structure prediction of biomolecular interactions with AlphaFold 3. Nature. 2024; 630(8016):493– 500.

Adalja A, Inglesby T. Broad-spectrum antiviral agents: a crucial pandemic tool. Expert review of Anti-infective Therapy. 2019; 17(7):467–470.

Adasme MF, Linnemann KL, Bolz SN, Kaiser F, Salentin S, Haupt VJ, Schroeder M. PLIP 2021: expanding the scope of the protein–ligand interaction profiler to DNA and RNA. Nucleic acids research. 2021; 49(W1):W530– W534.

Ahdritz G, Bouatta N, Floristean C, Kadyan S, Xia Q, Gerecke W, O’Donnell TJ, Berenberg D, Fisk I, Zanichelli N, Zhang B, Nowaczynski A, Wang B, Stepniewska-Dziubinska MM, Zhang S, Ojewole A, Guney ME, Biderman S, Watkins AM, Ra S, et al. OpenFold: Retraining AlphaFold2 yields new insights into its learning mechanisms and capacity for generalization. bioRxiv. 2022; https://www.biorxiv.org/content/10.1101/2022.11.20.517210, doi: 10.1101/2022.11.20.517210.

Altschul SF, Gish W, Miller W, Myers EW, Lipman DJ. Basic local alignment search tool. Journal of molecular biology. 1990; 215(3):403–410.

Anderson AC. The process of structure-based drug design. Chemistry & biology. 2003; 10(9):787–797.

ASAP Discovery Consortium, AI-driven structure-enabled antiviral platform (ASAP); 2023. Accessed: 2025-0421. https://asapdiscovery.org/.

Baek M, DiMaio F, Anishchenko I, Dauparas J, Ovchinnikov S, Lee GR, Wang J, Cong Q, Kinch LN, Schaeffer RD, et al. Accurate prediction of protein structures and interactions using a three-track neural network. Science. 2021; 373(6557):871–876.

Balcomb B, Marples P, Koekemoer L, Tomlinson C, Fearon D. Crystallisation of MERS-CoV Mpro in C2221 used for compound soaking. protocolsio. 2025 05; https://dx.doi.org/10.17504/protocols.io.8epv5rn84g1b/v2.

Balcomb B, Marples P, Koekemoer L, Tomlinson C, Fearon D. Crystallisation of SARS-CoV-2 MPro in P212121 for compound soaking V.2. protocolsio. 2025 05; https://dx.doi.org/10.17504/protocols.io.261ge5emyg47/v2.

Berman H, Henrick K, Nakamura H, Announcing the worldwide Protein Data Bank; 2003. doi: 10.1038/nsb1203-980.

Bloom JD, Neher RA. Fitness effects of mutations to SARS-CoV-2 proteins. Virus Evolution. 2023; 9(2):vead055.

Boby ML, Fearon D, Ferla M, Filep M, Koekemoer L, Robinson MC, Consortium‡ CM, Chodera JD, Lee AA, London N, et al. Open science discovery of potent noncovalent SARS-CoV-2 main protease inhibitors. Science. 2023; 382(6671):eabo7201.

Carvalho T, Krammer F, Iwasaki A. The first 12 months of COVID-19: a timeline of immunological insights. Nature Reviews Immunology. 2021; 21(4):245–256.

Chai Discovery. Chai-1: Decoding the molecular interactions of life. bioRxiv. 2024; https://www.biorxiv.org/content/early/2024/10/11/2024.10.10.615955, doi: 10.1101/2024.10.10.615955.

Cock PJ, Antao T, Chang JT, Chapman BA, Cox CJ, Dalke A, Friedberg I, Hamelryck T, Kauff F, Wilczynski B, et al. Biopython: freely available Python tools for computational molecular biology and bioinformatics. Bioinformatics. 2009; 25(11):1422.

Dai W, Zhang B, Jiang XM, Su H, Li J, Zhao Y, Xie X, Jin Z, Peng J, Liu F, et al. Structure-based design of antiviral drug candidates targeting the SARS-CoV-2 main protease. Science. 2020; 368(6497):1331–1335.

von Delft A, Hall MD, Kwong AD, Purcell LA, Saikatendu KS, Schmitz U, Tallarico JA, Lee AA. Accelerating antiviral drug discovery: lessons from COVID-19. Nature Reviews Drug Discovery. 2023; 22(7):585–603.

Eastman P, Galvelis R, Peláez RP, Abreu CR, Farr SE, Gallicchio E, Gorenko A, Henry MM, Hu F, Huang J, et al. OpenMM 8: molecular dynamics simulation with machine learning potentials. The Journal of Physical Chemistry B. 2023; 128(1):109–116.

Eberhardt J, Santos-Martins D, Tillack AF, Forli S. AutoDock Vina 1.2. 0: New docking methods, expanded force field, and python bindings. Journal of chemical information and modeling. 2021; 61(8):3891–3898.

Edwards A, Hartung IV. No shortcuts to SARS-CoV-2 antivirals. Science. 2021; 373(6554):488–489.

Elfiky AA. Anti-HCV, nucleotide inhibitors, repurposing against COVID-19. Life sciences. 2020; 248:117477.

Enard D, Cai L, Gwennap C, Petrov DA. Viruses are a dominant driver of protein adaptation in mammals. elife. 2016; 5:e12469.

Evans R, O’Neill M, Pritzel A, Antropova N, Senior A, Green T, Zidek A, Bates R, Blackwell S, Yim J, Ronneberger O, Bodenstein S, Zielinski M, Bridgland A, Potapenko A, Cowie A, Tunyasuvunakool K, Jain R, Clancy E, Kohli P, et al. Protein complex prediction with AlphaFold-Multimer. bioRxiv. 2021; doi: 10.1101/2021.10.04.463034v1.

Francoeur PG, Masuda T, Sunseri J, Jia A, Iovanisci RB, Snyder I, Koes DR. Three-dimensional convolutional neural networks and a cross-docked data set for structure-based drug design. Journal of chemical information and modeling. 2020; 60(9):4200–4215.

Geraghty RJ, Aliota MT, Bonnac LF. Broad-spectrum antiviral strategies and nucleoside analogues. Viruses. 2021; 13(4):667.

Gohlke H, Hendlich M, Klebe G. Knowledge-based scoring function to predict protein-ligand interactions. Journal of molecular biology. 2000; 295(2):337–356.

Goldin I. The Butterfly Defect: Why globalization creates systemic risks and what to do about it. Journal of Risk Management in Financial Institutions. 2014; 7(4):325–327.

Griffen EJ, Boulet P, Center AD, Moonshot C. Enabling equitable and affordable access to novel therapeutics for pandemic preparedness and response via creative intellectual property agreements. Wellcome Open Research. 2024; 9:374.

Guedes IA, Pereira FS, Dardenne LE. Empirical scoring functions for structure-based virtual screening: applications, critical aspects, and challenges. Frontiers in pharmacology. 2018; 9:1089.

Hamelryck T, Manderick B. PDB file parser and structure class implemented in Python. Bioinformatics. 2003; 19(17):2308–2310.

Jin Z, Du X, Xu Y, Deng Y, Liu M, Zhao Y, Zhang B, Li X, Zhang L, Peng C, et al. Structure of Mpro from SARS-CoV-2 and discovery of its inhibitors. Nature. 2020; 582(7811):289–293.

Judd EN, Gilchrist AR, Meyerson NR, Sawyer SL. Positive natural selection in primate genes of the type I interferon response. BMC Ecology and Evolution. 2021; 21(1):65.

Jumper J, Evans R, Pritzel A, Green T, Figurnov M, Ronneberger O, Tunyasuvunakool K, Bates R, Žídek A, Potapenko A, et al. Highly accurate protein structure prediction with AlphaFold. nature. 2021; 596(7873):583– 589.

Kabsch W. A solution for the best rotation to relate two sets of vectors. Foundations of Crystallography. 1976; 32(5):922–923.

Kelley BP, Brown SP, Warren GL, Muchmore SW. POSIT: flexible shape-guided docking for pose prediction. Journal of Chemical Information and Modeling. 2015; 55(8):1771–1780.

Lab V, PLIPify; 2021. Accessed: 2025-06-30. https://volkamerlab.org/projects/plipify/.

Lahav N, Haim Barr NL. MERS-CoV Mpro fluorescence dose response for antiviral testing. protocolsio. 2025 06; https://dx.doi.org/10.17504/protocols.io.eq2ly7r1rlx9/v5.

Lahav N, Haim Barr NL. SARS-CoV-2 Mpro fluorescence dose response. protocolsio. 2025 06; https://dx.doi.org/10.17504/protocols.io.81wgbye9nvpk/v5.

Leonard RA, Rao VN, Bartlett A, Froggatt HM, Luftig MA, Heaton BE, Heaton NS. A low-background, fluorescent assay to evaluate inhibitors of diverse viral proteases. Journal of Virology. 2023; 97(8):e00597–23.

Lin Z, Akin H, Rao R, Hie B, Zhu Z, Lu W, Smetanin N, Verkuil R, Kabeli O, Shmueli Y, et al. Evolutionary-scale prediction of atomic-level protein structure with a language model. Science. 2023; 379(6637):1123–1130.

Liu J, Wang R. Classification of current scoring functions. Journal of chemical information and modeling. 2015; 55(3):475–482.

Liu Y, Liang C, Xin L, Ren X, Tian L, Ju X, Li H, Wang Y, Zhao Q, Liu H, et al. The development of Coronavirus 3C-Like protease (3CLpro) inhibitors from 2010 to 2020. European journal of medicinal chemistry. 2020; 206:112711.

Madhav N, Oppenheim B, Gallivan M, Mulembakani P, Rubin E, Wolfe N. Pandemics: risks, impacts, and mitigation. Disease control priorities: improving health and reducing poverty 3rd edition. 2017;.

Mcgann MR, Almond HR, Nicholls A, Grant JA, Brown FK. Gaussian docking functions. Biopolymers: Original Research on Biomolecules. 2003; 68(1):76–90.

McNutt AT, Francoeur P, Aggarwal R, Masuda T, Meli R, Ragoza M, Sunseri J, Koes DR. GNINA 1.0: molecular docking with deep learning. Journal of cheminformatics. 2021; 13(1):43.

McNutt AT, Li Y, Meli R, Aggarwal R, Koes DR. GNINA 1.3: the next increment in molecular docking with deep learning. Journal of Cheminformatics. 2025; 17(1):28.

Mey AS, Allen BK, Macdonald HEB, Chodera JD, Hahn DF, Kuhn M, Michel J, Mobley DL, Naden LN, Prasad S, et al. Best practices for alchemical free energy calculations [article v1. 0]. Living journal of computational molecular science. 2020; 2(1):18378.

Mirdita M, Schütze K, Moriwaki Y, Heo L, Ovchinnikov S, Steinegger M. ColabFold: making protein folding accessible to all. Nature methods. 2022; 19(6):679–682.

Muegge I, Martin YC. A general and fast scoring function for proteinligand interactions: a simplified potential approach. Journal of medicinal chemistry. 1999; 42(5):791–804.

Naqvi AAT, Fatima K, Mohammad T, Fatima U, Singh IK, Singh A, Atif SM, Hariprasad G, Hasan GM, Hassan MI. Insights into SARS-CoV-2 genome, structure, evolution, pathogenesis and therapies: Structural genomics approach. Biochimica et Biophysica Acta (BBA)-Molecular Basis of Disease. 2020; 1866(10):165878.

Noske GD, de Souza Silva E, de Godoy MO, Dolci I, Fernandes RS, Guido RVC, Sjö P, Oliva G, Godoy AS. Structural basis of nirmatrelvir and ensitrelvir activity against naturally occurring polymorphisms of the SARS-CoV-2 main protease. Journal of Biological Chemistry. 2023; 299(3).

OpenEye, Cadence Molecular Sciences. OpenEye Toolkits 2024.2.1. OpenEye, Cadence Molecular Sciences, Santa Fe, NM; 2024, http://www.eyesopen.com.

OpenEye, Cadence Molecular Sciences, Inc. OEDOCKING 4.3.2.1. OpenEye, Cadence Molecular Sciences, Inc., Santa Fe, NM; 2025, http://www.eyesopen.com.

Passaro S, Corso G, Wohlwend J, Reveiz M, Thaler S, Somnath VR, Getz N, Portnoi T, Roy J, Stark H, Kwabi-Addo D, Beaini D, Jaakkola T, Barzilay R. Boltz-2: Towards Accurate and Efficient Binding Affinity Prediction. bioRxiv. 2025; doi: 10.1101/2025.06.14.659707.

Salentin S, Schreiber S, Haupt VJ, Adasme MF, Schroeder M. PLIP: fully automated protein–ligand interaction profiler. Nucleic acids research. 2015; 43(W1):W443–W447.

Sayers EW, Beck J, Bolton EE, Bourexis D, Brister JR, Canese K, Comeau DC, Funk K, Kim S, Klimke W, et al. Database resources of the national center for biotechnology information. Nucleic acids research. 2021; 49(D1):D10–D17.

Schake P, Dishnica K, Kaiser F, Leberecht C, Haupt VJ, Schroeder M. An interaction-based drug discovery screen explains known SARS-CoV-2 inhibitors and predicts new compound scaffolds. Scientific Reports. 2023; 13(1):9204.

Schrödinger, LLC. Schrödinger Release 2025-2: Maestro. Schrödinger, LLC, New York, NY; 2025, https://www.schrodinger.com/.

Schuler GD, Epstein JA, Ohkawa H, Kans JA. [10] Entrez: Molecular biology database and retrieval system. In: Methods in enzymology, vol. 266 Elsevier; 1996.p. 141–162.

Trott O, Olson AJ. AutoDock Vina: improving the speed and accuracy of docking with a new scoring function, efficient optimization, and multithreading. Journal of computational chemistry. 2010; 31(2):455–461.

Tummino TA, Rezelj VV, Fischer B, Fischer A, O’meara MJ, Monel B, Vallet T, White KM, Zhang Z, Alon A, et al. Drug-induced phospholipidosis confounds drug repurposing for SARS-CoV-2. Science. 2021; 373(6554):541– 547.

Vidal J. Destroyed habitat creates the perfect conditions for coronavirus to emerge. Scientific American. 2020; 18(03):2020.

Wang F, Chen C, Wang Z, Han X, Shi P, Zhou K, Liu X, Xiao Y, Cai Y, Huang J, et al. The structure of the porcine deltacoronavirus main protease reveals a conserved target for the design of antivirals. Viruses. 2022; 14(3):486.

Wang W, Zhao H, Han GZ. Host-virus arms races drive elevated adaptive evolution in viral receptors. Journal of virology. 2020; 94(16):10–1128.

Wohlwend J, Corso G, Passaro S, Getz N, Reveiz M, Leidal K, Swiderski W, Atkinson L, Portnoi T, Chinn I, Silterra J, Jaakkola T, Barzilay R. Boltz-1: Democratizing Biomolecular Interaction Modeling. bioRxiv. 2024; doi: 10.1101/2024.11.19.624167.

Yang H, Xie W, Xue X, Yang K, Ma J, Liang W, Zhao Q, Zhou Z, Pei D, Ziebuhr J, et al. Design of wide-spectrum inhibitors targeting coronavirus main proteases. PLoS biology. 2005; 3(10):e324.

